# Unraveling IGK Locus in Dog Breeds: IMGT® New Insights into Canine Immunogenetics

**DOI:** 10.1101/2025.11.10.687733

**Authors:** Taismara Kustro Garnica, Ariadni Papadaki, Maria Georga, Guilhem Zeitoun, Joumana Jabado-Michaloud, Géraldine Folch, Véronique Giudicelli, Pablo Gonzales, Jéssika Cristina Chagas Lesbon, Talita Gabriela Luna Alves, Heidge Fukumasu, Sofia Kossida

## Abstract

Over millennia, the selective breeding of dogs (*Canis lupus familiaris*) has generated remarkable genetic diversity among breeds, highlighting the need for comprehensive genomic and immunogenetic studies. This research provides detailed immunoglobulin kappa light chain locus (IGK) analysis across multiple dog breeds. It aims to uncover breed-specific genetic variations and their implications for immunology and veterinary medicine. The primary objectives were to do the biocuration of the IGK locus in nine canine genome assemblies, investigate structural variations, polymorphisms, and gene diversity, and to enrich the IMGT® database with comprehensive IGK data from diverse breeds, creating a more inclusive genetic resource. Our extensive annotation of breeds, including the Bernese Mountain Dog, Boxer, Cairn Terrier, Labrador Retriever, Great Dane, Basenji, and German Shepherd, identified 40 genes and 97 alleles, revealing both conserved genes and unique variants across these breeds, with in silico validation through Sanger sequencing. Notably, we analyzed discrepancies in the first reference assembly from the Boxer breed (Canfam3.1), highlighting potential errors in assembly, challenges in gene and allele nomenclature, and a low-density region within the canine IGK locus. This study not only refines the understanding of IGK locus diversity but also contributes to the IMGT® databases, advancing future research on immunogenetic variability, somatic mutations, and immune response dynamics in canine health and disease.

## 1. Introduction

Immunoglobulins are composed of two identical heavy chains (IGH) and two identical light chains, either kappa (IGK) or lambda (IGL), connected by disulfide bonds. The heavy chains are encoded by variable (V), diversity (D), joining (J), and constant (C) genes, while the light chains are encoded by V, J, and C genes (Lefranc & Lefranc, 2022). The IGK light chain exhibits less diversity than the IGH across species (Collins & Watson, 2018). However, the IGK germline diversity in dog breeds is still largely uncharacterized, which can be attributed to the additional complexity introduced by high-frequency repeat regions, which makes the annotation of IG genes particularly challenging (Cullen et al., 2022; Sirupurapu et al., 2022). Research on immunoglobulin (IG) and T-cell receptor (TR) genes across dog breeds is crucial for building a comprehensive genetic database to advance veterinary medicine. Such research is essential for understanding breed-specific predispositions to immune-related diseases, including lymphoma—a hematological malignancy of lymphocytes—that is well-documented in breeds like Doberman Pinschers, Rottweilers, Boxers, and Bernese Mountain Dogs (Comazzi et al., 2018; Hédan et al., 2021). Moreover, identifying allelic variants and polymorphisms associated with conditions such as chronic lymphocytic leukemia (CLL) and diffuse large B-cell lymphoma (DLBCL) offers valuable insights into disease mechanisms (Avery, 2020; Rout et al., 2018). Despite these advances, the genetic diversity of immunoglobulins (IG), which function as B-cell antigen receptors, remains underexplored across different breeds, highlighting a critical gap in our understanding of canine immune genetics.

The International Immunogenetics Information System (IMGT®), established in 1989, serves as the global standard for immunogenetics, providing databases and tools critical for investigating the adaptive immune response (Manso et al., 2022). According to IMGT®, the canine IGK locus, the smallest of the IG loci, is located on chromosome 17, spanning 317-349 kb and comprising 31 genes (25 V, 5 J, 1 C) based on two reference assemblies: Basenji (Basenji_breed-1.0) and Boxer (CanFam3.1), corresponding to IMGT/LIGM-DB accession numbers IMGT000067 and IMGT000002, respectively (Martin et al., 2018). However, discrepancies in the V-J-C-CLUSTER and locus orientation between these references require further investigation for accurate biocuration of novel canine assemblies. To address these challenges, this study has four main objectives: (1) conduct a detailed comparative analysis to verify the orientation and gene positions in the dog IGK locus reference assemblies; (2) perform extensive biocuration of the IGK locus across nine canine genome assemblies to uncover structural variations, polymorphisms, and gene and allele diversity; (3) enhance the IMGT® databases, tools, reference directories, and web resources by integrating data from multiple dog breeds, thereby creating a more comprehensive and diverse genetic repository; and (4) confirm the IGKV allelic variation through Sanger sequencing.

## 2. Results

### 2.1. Comprehensive annotation of the IGK locus across dog breeds

#### 2.1.1. Quality evaluation of IGK locus assemblies

During the quality assessment of the loci, it was observed that three assemblies (33.3%) presented small gaps in the IGK locus. The Bernese Mountain Dog (OD_1.0), Labrador Retriever (Yella_v2), and Great Dane (UMICH_Zoey_3.1) with variable sizes from 99 to 200 bp (Supplemental_Table_S1). However, the genomic structure (V-J-C-CLUSTER) and orientation of the IGK locus were not affected, and they passed the quality control of IMGT.

#### 2.1.2. Localization and description of the IGK locus

The biocuration of dog IGK locus followed the IMGT standards and comprised seven distinct dog breeds: Basenji, Bernese Mountain Dog, Boxer, Cairn Terrier, German Shepherd, Great Dane, and Labrador Retriever. Accounting with five assemblies from female dogs and four from male dogs. The IGK locus in dogs had a variable length, ranging from 235 kb to 369 kb across the nine assemblies, with the longest locus belonging to the Bernese Mountain Dog assembly (OD_1.0) and the shortest to the Boxer (Dog10K_Boxer_Tasha) (Table 1). The highest number of V genes was found in both Bernese Mountain Dog assemblies (BD_1.0 and OD_1.0) and the lowest was found in Boxer (Dog10K_Boxer_Tasha). However, the number of J and C genes was conserved among the assemblies (Table 1).

**Table 1.** Mainly data from the IGK locus biocuration from 9 dog assemblies.

| IMGT/LIGM-DB | Assembly (breed) | V | J | C | Total | Length (kb) |
| --- | --- | --- | --- | --- | --- | --- |
| IMGT000184 | ASM864105v1 (German Shepherd) | 21 | 5 | 1 | 26 | 264 |
| BK066061* | BD_1.0 (Bernese Mountain Dog) | 29 | 5 | 1 | 35 | 358 |
| IMGT000179 | CA611_1.0 (Cairn Terrier) | 24 | 5 | 1 | 30 | 298 |
| IMGT000185 | Dog10K_Boxer_Tasha (Boxer) | 19 | 5 | 1 | 25 | 235 |
| BK066062* | OD_1.0 (Bernese Mountain Dog) | 29 | 5 | 1 | 35 | 369 |
| IMGT000181 | ROS_Cfam_1.0 (Labrador Retriever) | 21 | 5 | 1 | 27 | 274 |
| IMGT000183 | UMICH_Zoey_3.1 (Great Dane) | 25 | 5 | 1 | 31 | 306 |
| IMGT000180 | UNSW_CanFamBas_1.2 (Basenji) | 23 | 5 | 1 | 29 | 285 |
| IMGT000182 | Yella_v2 (Labrador Retriever) | 22 | 5 | 1 | 28 | 279 |

\*Annotations available on TPA (third-party annotation, NCBI) with the same accession number as IMGT/LIGM-DB.

A holistic IGK locus map was created to integrate all data obtained from the annotated assemblies (gene positions, allele function, V-J-C-CLUSTER, and locus orientation) (Figure 1). The IGK locus was found forward-oriented (FWD) on chromosome 17, for all dog assemblies. The locus extends from 10 kb upstream, IGKV9-24, to 10 kb downstream, IGKC. The two IMGT flanking genes, referred to as “IMGT bornes”, were identified through BLAST alignment and annotated in all assemblies to define the boundaries of the IGK locus. According to the IGK holistic map for dogs, the 5’ end IMGT flanking gene, “IMGT borne”, PAX8 (Gene ID: 403927) is 137 kb upstream of IGKV9-24, the most 5’ gene in the locus, and the 3’ end IMGT flanking gene, “IMGT borne”, RPIA (Gene ID: 100856455) is 35 kb downstream of IGKC, the most 3’ gene in the locus. The IGK locus consists predominantly of a cluster of 34 V genes (V-CLUSTER) and a smaller cluster containing 5 J genes and 1 C gene (J-C-CLUSTER). Five genes were oriented in the reverse direction in the locus — IGKV4-15, IGKV(I)-14, IGKV3-3, IGKV7-2, and IGKV4-1— across all assemblies.

**Figure 1.**
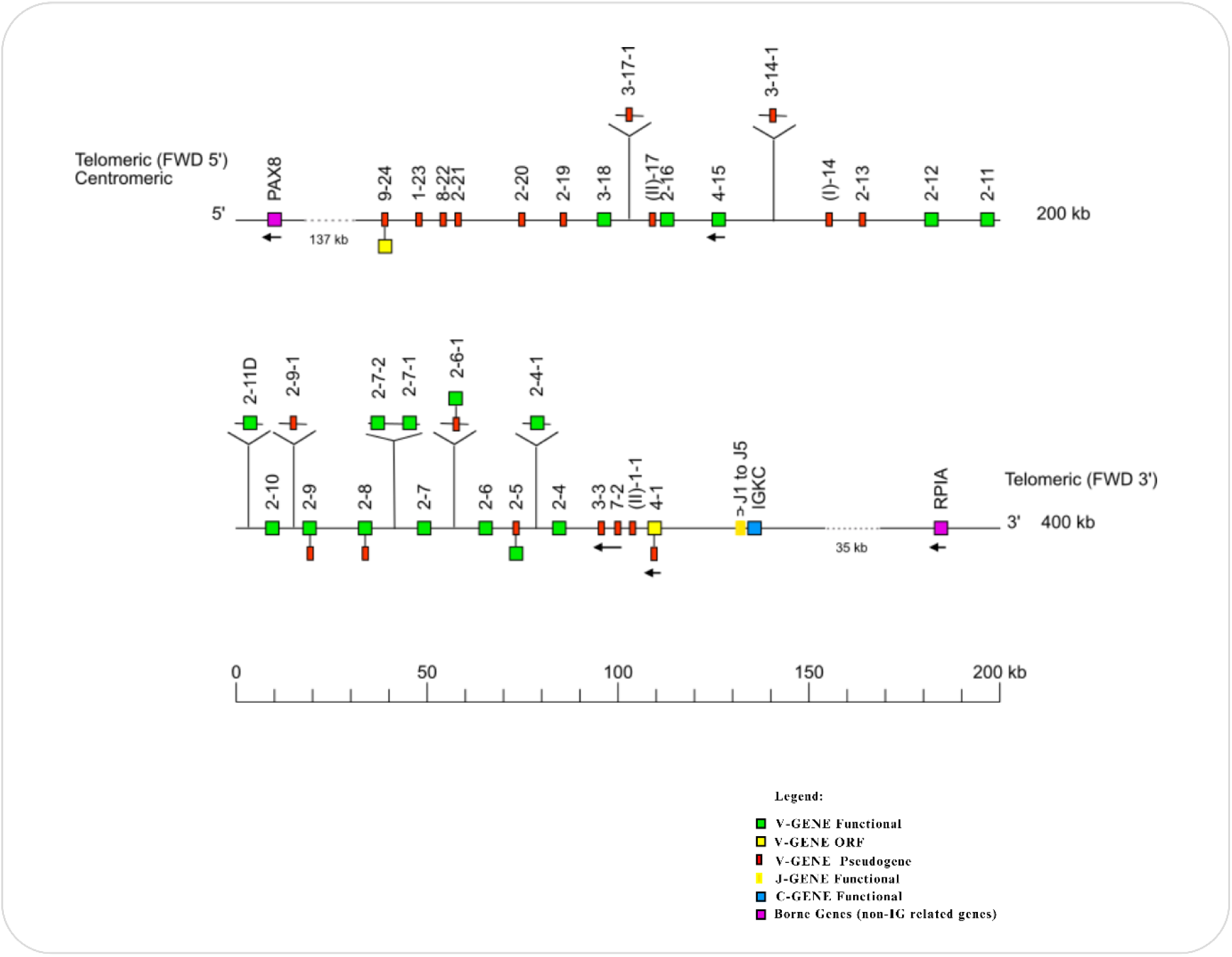
Holistic IMGT reference IGK map for dogs (*Canis lupus familiaris*). The dog IGK locus is on chromosome 17, and the orientation is forward (FWD). The colors are according to the IMGT® color menu for genes^6^ [^6^https://www.imgt.org/IMGTScientificChart/RepresentationRules/colormenu.php#LOCUS]. The boxes representing the genes are not scaled, and the exons are not shown. IGK gene names follow the IMGT nomenclature (Lefranc, 2014). The dog’s IGKV genes belong to nine subgroups IGKV1, IGKV2, IGKV3, IGKV4, IGKV7, IGKV8, IGKV9 as well as IGKV(I) and IGKV(II), belonging to clans^7^ [^7^*A clan is a set of subgroups, eventually from different species, which appear related to phylogenetic trees (IGHV, IGKV, IGLV clans). The ’clan’ concept is part of the ’CLASSIFICATION’ concept of IMGT-ONTOLOGY*.] (Giudicelli & Lefranc, 1999). The dots represent the distance between the borne genes (PAX8 and RPIA), the upstream gene (IGKV9-24), and the downstream gene (IGKC) in the locus. Single arrows show genes whose polarity is opposite to that of the J-C-CLUSTER comprising IGKJ (J1 to J5)-IGKC. The 8 highlighted genes above the central axis represent genes not found in all assemblies. The central axis represents the functionality of the alleles found in the reference genomes (CanFam3.1 and Basenji_breed-1.1)—reproduction authorized by IMGT®, the international ImMunoGeneTics information system, available on the IMGT® website at https://www.imgt.org/IMGTrepertoire/index.php?section=LocusGenes&repertoire=locus&species=dog&group=IGK&assembly=Holistic_IMGT_reference.

#### 2.1.3. Annotation of IGKV gene cluster

The biocuration of dog assemblies identified 34 IGKV genes (Figure 1). In dogs, the IGKV genes are categorized into nine distinct subgroups (IGKV1, IGKV2, IGKV3, IGKV4, IGKV7, IGKV8, and IGKV9) and two clans IGKV(I) and IGKV(II), according to the IMGT-ONTOLOGY and their sequence correspondence with the IGK human subgroups (Figure 1). Among these, the IGKV2 subgroup was the most representative, encompassing 21 genes constituting 61.76% of the total V genes, followed by IGKV3 with 4 genes (11.76%).

Seven new genes were discovered during the annotation of nine dog assemblies (IGKV3-17-1, IGKV2-11D, IGKV2-9-1, IGKV2-7-2, IGKV2-7-1, IGKV2-6-1, and IGKV2-4-1), accounting for a discovery rate of 17.5%. However, these new genes were uncommon for all the assemblies (Table 3). The IGKV2-7-1 gene was found in two distinct breed assemblies from Bernese Mountain Dog (BD_1.0) and Cairn Terrier (CA611_1.0). The IGKV2-6-1 gene was found in four assemblies from Bernese Mountain Dog (BD_1.0 and OD_1.0), Cairn Terrier (CA611_1.0), and Basenji (UNSW_CanFamBas_1.2). Finally, the IGKV2-4-1 gene was found in two assemblies from the Bernese Mountain Dog (BD_1.0 and OD_1.0). Nevertheless, four genes were found exclusively in one assembly. The IGKV3-17-1 and IGKV2-11D genes were identified only in the Great Dane assembly (UMICH_Zoey_3.1), and the duplicated gene was found in this assembly at the contig level. The IGKV2-9-1 gene was unique to the Bernese Mountain Dog assembly (OD_1.0), while the IGKV2-7-2 gene was exclusively in the Bernese Mountain Dog assembly (BD_1.0). According to our results, most new IGKV genes (5 out of 7, 71.42%) were identified in the Bernese Mountain Dog breed assemblies (Table 3).

**Table 2.** Dogs (*Canis lupus familiaris*) IGKV genes per subgroup.

| IGKV Subgroup | Number of genes | Freq. (%) |
| --- | --- | --- |
| IGKV1 | 1 | 2.94 |
| IGKV2 | 21 | 61.76 |
| IGKV3 | 4 | 11.76 |
| IGKV4 | 2 | 5.88 |
| **IGKV7** | 1 | 2.94 |
| **IGKV8** | 1 | 2.94 |
| **IGKV9** | 1 | 2.94 |
| **IGKV(I)** | 1 | 2.94 |
| **IGKV(II)** | 2 | 5.88 |
| **Total** | 34 | 100 |

**Table 3.** New IGKV genes per assembly.

| **New IGKV genes** | **Breed (Assembly)** |
| --- | --- |
| **IGKV3-17-1** | Great Dane (UMICH_Zoey_3.1) |
| **IGKV2-11D** | Great Dane (UMICH_Zoey_3.1) |
| **IGKV2-9-1** | Bernese Mountain Dog (OD_1.0) |
| **IGKV2-7-2** | Bernese Mountain Dog (BD_1.0) |
| **IGKV2-7-1** | Bernese Mountain Dog (BD_1.0)<br>Cairn Terrier (CA611_1.0) |
| **IGKV2-6-1** | Bernese Mountain Dog (BD_1.0)<br>Bernese Mountain Dog (OD_1.0)<br>Cairn Terrier (CA611_1.0)<br>Basenji<br>(UNSW_CanFamBas_1.2). |
| **IGKV2-4-1** | Bernese Mountain Dog (BD_1.0)<br>Bernese Mountain Dog (OD_1.0) |

#### 2.1.4. Annotation of IGKJ and IGKC gene clusters

All nine canine assemblies conserved five IGKJ genes (J-CLUSTER), each belonging to a different set (J1, J2, J3, J4, and J5) and one IGKC gene (C-CLUSTER).

### 2.2. Insights into canine immunogenetic diversity

#### 2.2.1. Patterns of gene conservation across dog breeds

The overall conservation of IGK genes across dog assemblies was 50%, with 20 out of 40 IGKV, IGKJ, and IGKC genes being shared. This analysis included genes identified in all annotated dog assemblies available on the IMGT website, comprising two IMGT reference assemblies and nine newly annotated assemblies (Figure 2). Conversely, the remaining 50% of IGK genes, all from the IGKV group, exhibited variability across the dog assemblies. Notably, a unique gene, IGKV2S13, identified in the IMGT reference Boxer assembly (CanFam3.1), was absent from all newly annotated dog assemblies in this study.

**Figure 2.**
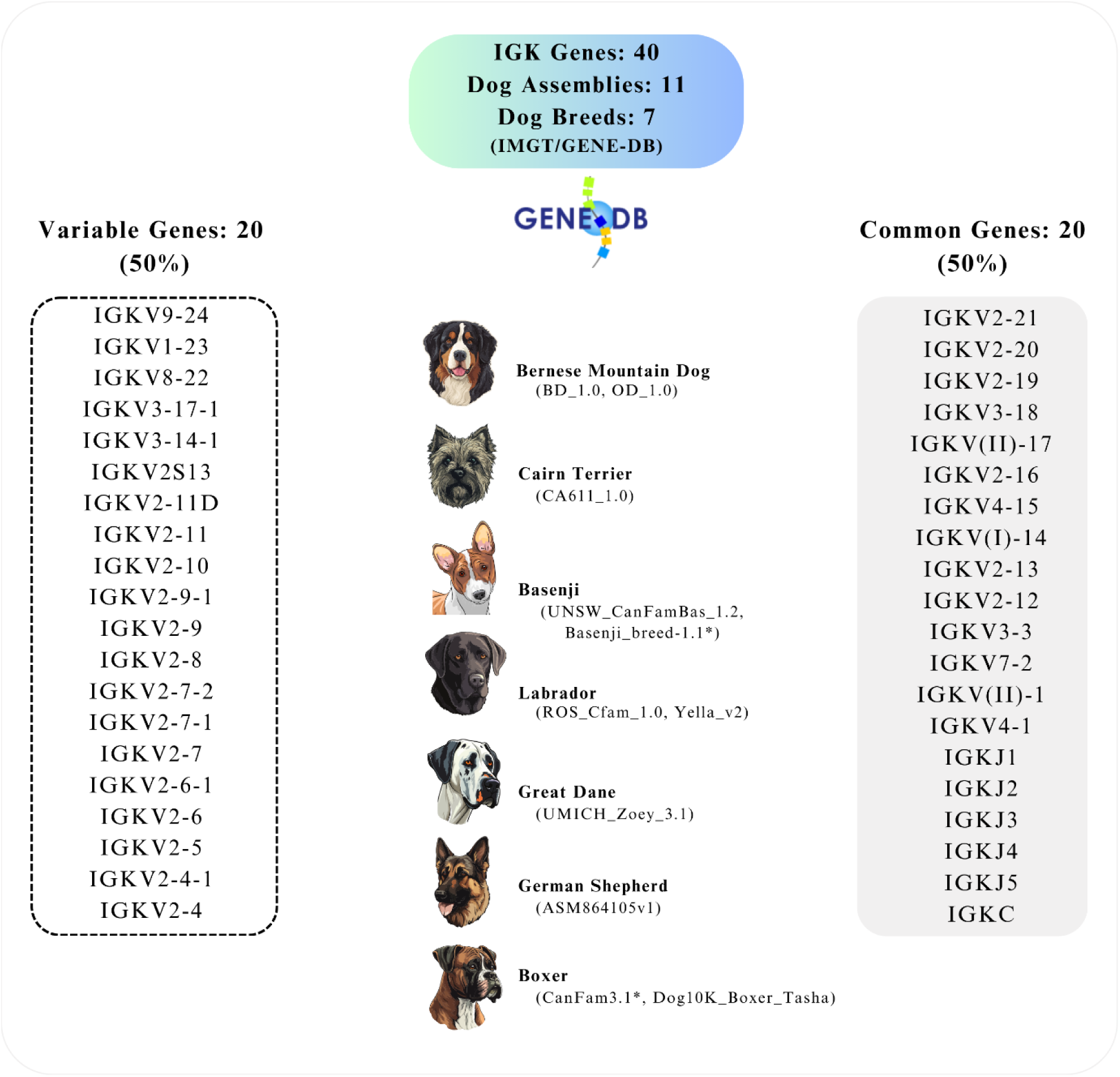
Representation of the IGK locus genes in dogs (*Canis lupus familiaris*) that are either common (100% frequency) or variable (<100% frequency) across 11 annotated dog breed assemblies (2 IMGT reference and 9 new assemblies) available on IMGT/GENE-DB (https://www.imgt.org/genedb/). Notably, the conservation rate of IGK genes within this locus is 50% across all breeds (https://imgt.org/IMGTrepertoire/LocusGenes/locusdesc/assembly_compare.php?species=Canis+lupus+familiaris&locus=IGK).

#### 2.2.2. Discovery of new alleles

During this study, it was possible to characterize 97 IGK alleles, most of which belonged to the variable genes (IGKV). No new alleles were identified for the IGKJ and IGKC genes beyond those previously characterized in the IMGT reference assemblies (CanFam3.1 and Basenji_breed-1.1). A single allele was found for each IGKJ gene (IGKJ1*01, IGKJ2*01, IGKJ3*01, IGKJ4*01, and IGKJ5*01), and two alleles were identified for the IGKC constant gene (IGKC*01 and IGKC*02).

90 alleles were identified across 34 IGKV genes (Figure 3). Of these, 29 genes were polymorphic, while only 5 genes exhibited the same allele across all dog assemblies: IGKV1-23, IGKV2-19, IGKV4-15, IGKV(I)-14, and IGKV(II)-1-1 (Figure 4). The new alleles were defined based on the percentage of identity in the V-REGION ranging from 96% to 99%. The newly identified alleles for each assembly are highlighted in yellow, along with their respective identity percentages, and can be found in the supplemental material (Supplemental_Table_S2). The majority of dog IGKV genes belong to the IGKV2 subgroup (Figure 3 and Table 2). The varying population sizes within a given immunoglobulin V subgroup can be explained by factors such as genetic evolution, selective pressures, and recombination preferences observed in other species (Mage et al., 2016).

**Figure 3.**
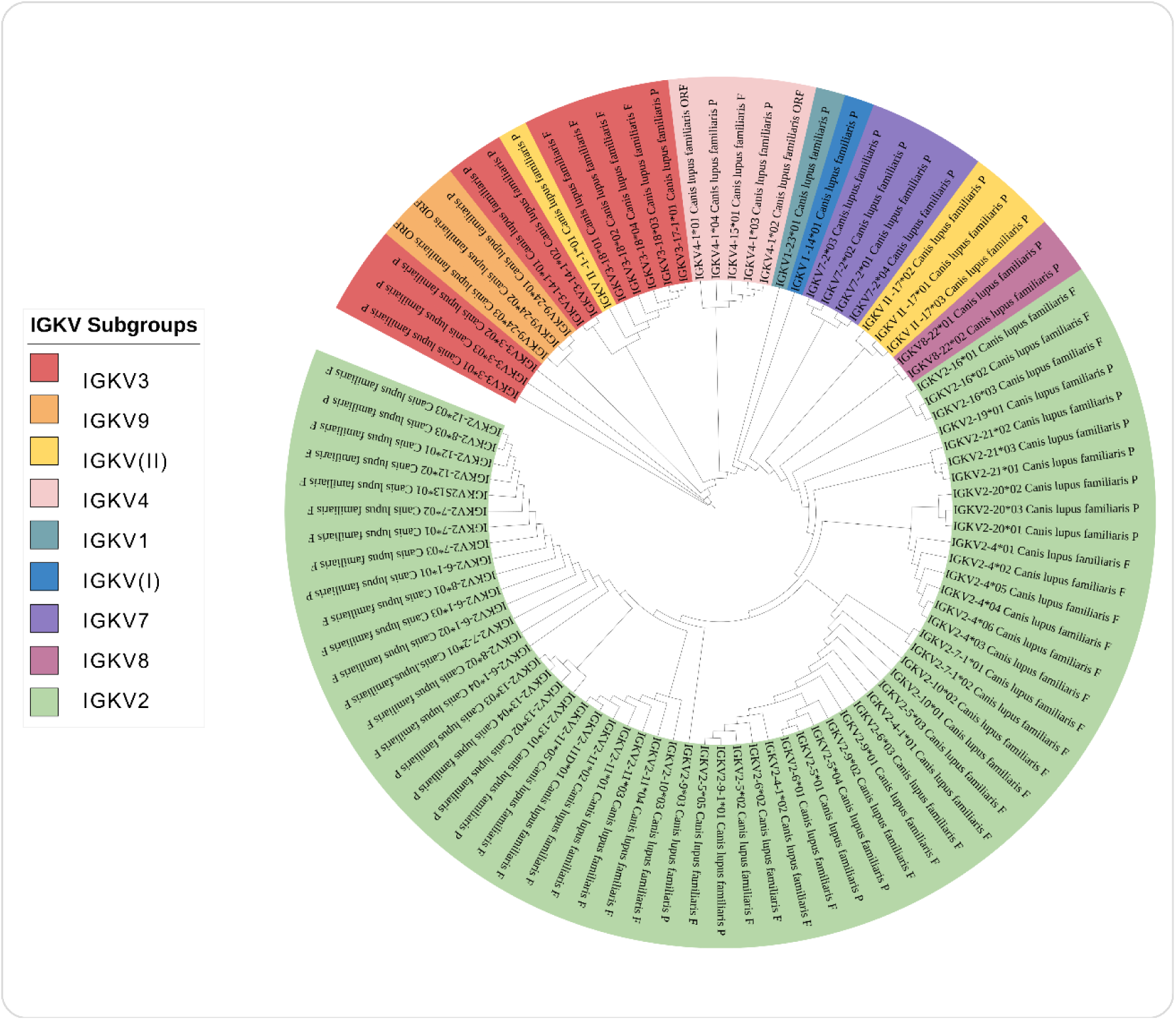
A comprehensive phylogenetic tree displaying 90 IGKV alleles of the dog (*Canis lupus familiaris*) categorized by their respective IGKV subgroups, based on V-region sequence comparison. The tree was constructed using NGPhylogeny.fr (Lemoine et al., 2019), employing MAFFT version 7 (Rozewicki et al., 2019) for multiple sequence alignment, PhyML (Guindon et al., 2010) for maximum likelihood phylogenetic inference, and iTOL v6 (Letunic & Bork, 2024) for visualization.

**Figure 4.**
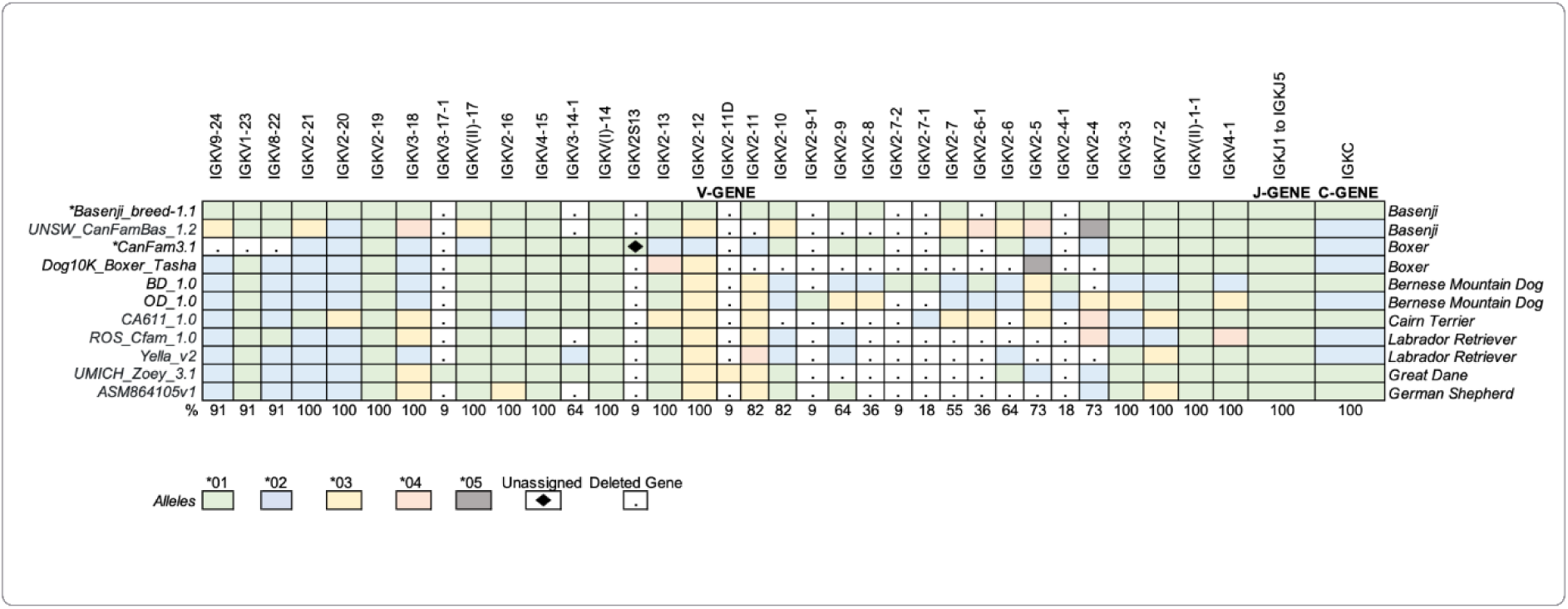
The heatmap displays the allelic polymorphism at the gDNA level of the dog (*Canis lupus familiaris*) IGK locus, based on 11 genome assemblies from distinct dog breeds. Each row represents a locus from a specific assembly/breed, and the columns represent each IGK gene in their respective order within the locus (V-J-C-CLUSTER). Each color corresponds to different alleles (from *01 to *06). Blank squares indicate missing genes in each assembly (unassigned). Squares with a diamond shape represent a specific allele, IGKV2S13*01 found only in one assembly (CanFam3.1) with provisional nomenclature. Gene frequencies are presented below.

Our analysis found a common “low coverage region” in the V-CLUSTER of all the assemblies. The region between IGKV2-9 and IGKV2-4 had fewer genes than the locus’s upstream and downstream regions. Interestingly, the genes at low frequency in this region had a high diversity of alleles, as we can see from IGKV2-5 and IGKV2-4 (Figure 4).

Considering the functionality of the IGKV alleles, most of them are functional. The IGKV2 subgroup exhibited the highest allele count, with 60 alleles (66.6%), predominantly functional, including 43 F and 17 P alleles. The V3 subgroup contained the second-largest number of alleles, comprising 6 functional alleles (F) and 4 pseudogenes (P). Subgroups V4 and V9 contained two ORFs. Subgroup V1 and clan (I) presented a unique allele, both P. In total, the analysis identified 48 F (53.3%), 38 P (42.2%), and 4 ORF (4.5%) IGKV alleles (Table 4).

**Table 4.** The dog (*Canis lupus familiaris*) IGK alleles frequency per subgroup and their functionality.

| **Subgroups** | **n** | **Freq. (%)** | **Functionality** |
| --- | --- | --- | --- |
| **V2** | 60 | 66.6 | 43 (F), 17 (P) |
| **V3** | 10 | 11.1 | 4 (F), 6 (P) |
| **V4** | 5 | 5.5 | 1 (F), 2 (ORF), 2 (P) |
| **V7** | 4 | 4.4 | 4 (P) |
| **V(II)** | 4 | 4.4 | 4 (P) |
| **V9** | 3 | 3.3 | 2 (ORF), 1 (P) |
| **V8** | 2 | 2.2 | 2 (P) |
| **V1** | 1 | 1.1 | 1 (P) |
| **V(I)** | 1 | 1.1 | 1 (P) |
| **Total** | 90 | 100 | 48 (F), 4 (ORF), 38 (P) |

### 2.3. Validation of new alleles by Sanger sequencing

Nine PCR products were successfully sequenced using the Sanger method with the seven designed primers. Among them, two out of seven primers yielded sequences showing 100% alignment identity with IGKV2 alleles deposited in the IMGT/GENE-DB database. Specifically, Primer 6 (FWD) amplified and confirmed the allele IGKV2-12*03 (F), while Primer 12 (FWD) identified the allele IGKV2-40*4 (F), confirming their existence and homozygosity in the analyzed samples (Figure 5).

**Figure 5.**
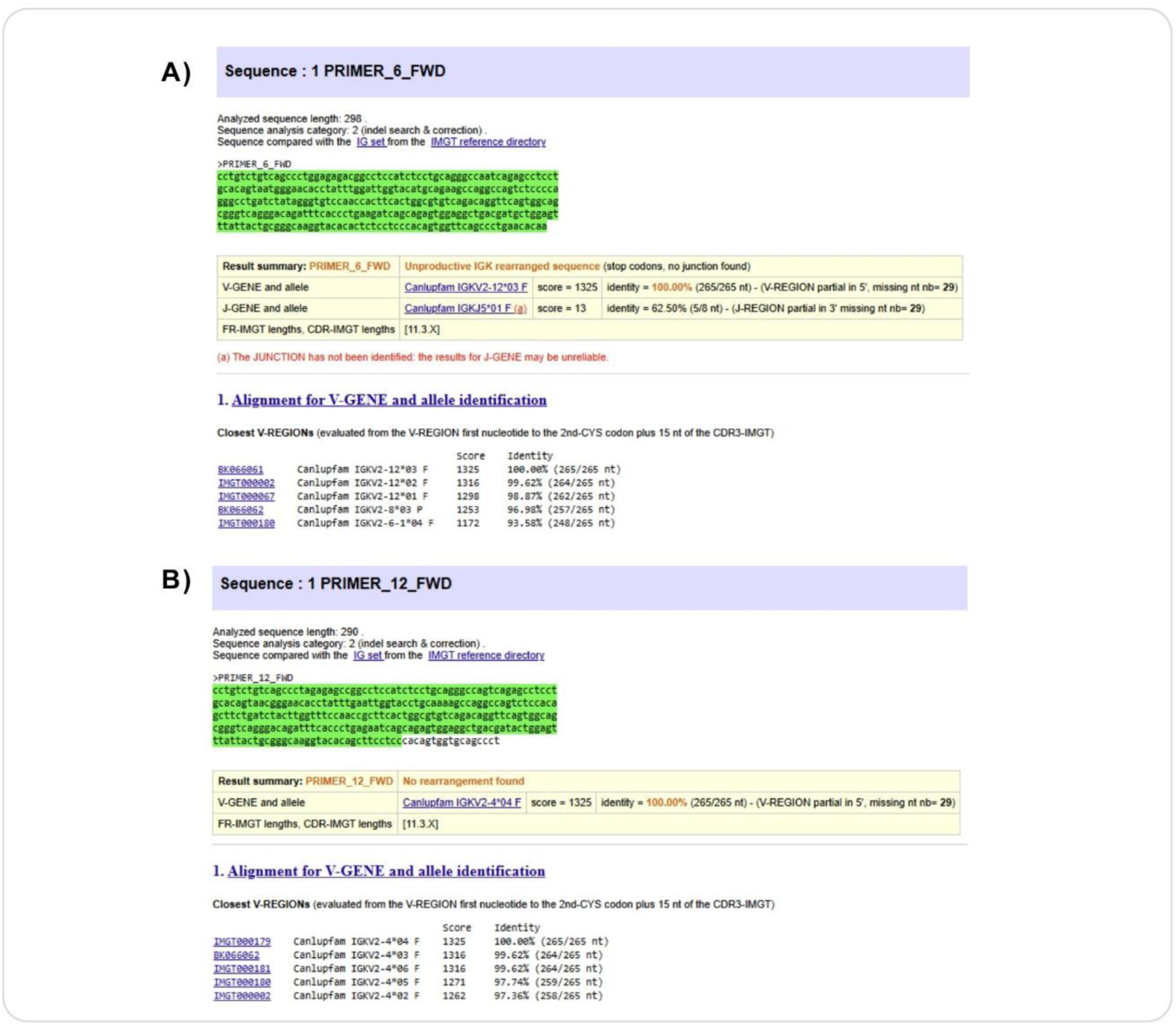
Validation of two annotated IGKV2 alleles using IMGT/V-QUEST analysis. (A) Alignment results confirming the allele IGKV2-12*03 obtained with forward primer 6; (B) alignment results confirming the allele IGKV2-4*04 obtained with forward primer 12.

The remaining five primers produced sequences with alignment identities ranging from 92.08% to 97.69%, suggesting the presence of heterozygous loci or allelic variants not yet represented in the reference database (Supplemental Table S4).

### 2.4. Challenges and limitations

#### 2.4.1. Locus annotation discrepancies in IMGT references

Before initiating annotations for the new dog assemblies, a thorough analysis of the IMGT reference sequences for the dog’s IGK locus was conducted. This step was necessary due to discrepancies in locus orientation, which could affect the accurate identification and extraction of this locus in subsequent assemblies. For the *Canis lupus familiaris* IGK locus, two reference assemblies from distinct breeds are available in the IMGT® database: the Boxer (CanFam3.1; accession number IMGT000002) and the Basenji (Basenji_breed-1.1; accession number IMGT000067), both listed in the IMGT/LIGM-DB.

The IGK locus in dogs is located on chromosome 17, with a reverse (REV) orientation in the Boxer (CanFam3.1), and a forward (FWD) orientation in the Basenji (Figure 5, A and B). In the Boxer (CanFam3.1) assembly, the position of the V-J-CLUSTER reveals unusual organization, with the J-C-CLUSTER positioned centrally within the IGK locus rather than at the expected 3’ end. In contrast, the Basenji assembly displays an IGK locus organization more consistent with other veterinary species, such as the cat, goat, sheep, horse, bovine, and pig, as represented in IMGT® > Repertoire > Locus representations (https://www.imgt.org/IMGTrepertoire/LocusGenes/#B).

The length of the IGK locus was 349 kb in the Boxer (CanFam3.1), assembly and 317 kb in the Basenji (Basenji_breed-1.1) assembly, showing two gaps in the Boxer, in the region of the IGK locus on chromosome 17. The smallest gap was 226 bp, between the positions 227,525 and 227,750 bp, while the largest gap was 950 bp, between the positions 337,118 and 338,067 bp. The 950 bp size gap was positioned between the IGKC*02 and IGKV2-12*02 genes, and the 226 bp gap occurred within the IGKV3-18 gene. Although for the IGK locus in Basenji (Basenji_breed-1.1), no gap was found (Figure 6, A and B). The flanking genes, referred to as ’IMGT bornes’, for the IGK locus were present in both reference assemblies, though the position of RPIA differed notably. In the Basenji assembly (Basenji_breed-1.1), the 5’ IMGT flanking gene, PAX8 (PAX8, Gene ID: 701906), is located 137 kb upstream of IGKV9-24, the most 5’ gene in the locus, while the 3’ IMGT flanking gene, RPIA (RPIA, Gene ID: 699694), is positioned 35 kb downstream of IGKC. In the Boxer reference (CanFam3.1), PAX8 is 161 kb upstream of IGKV2-11, the most 5’ gene in the locus, and RPIA is situated between the IGKC and IGKV2-12 genes (Figure 6, C).

**Figure 6.**
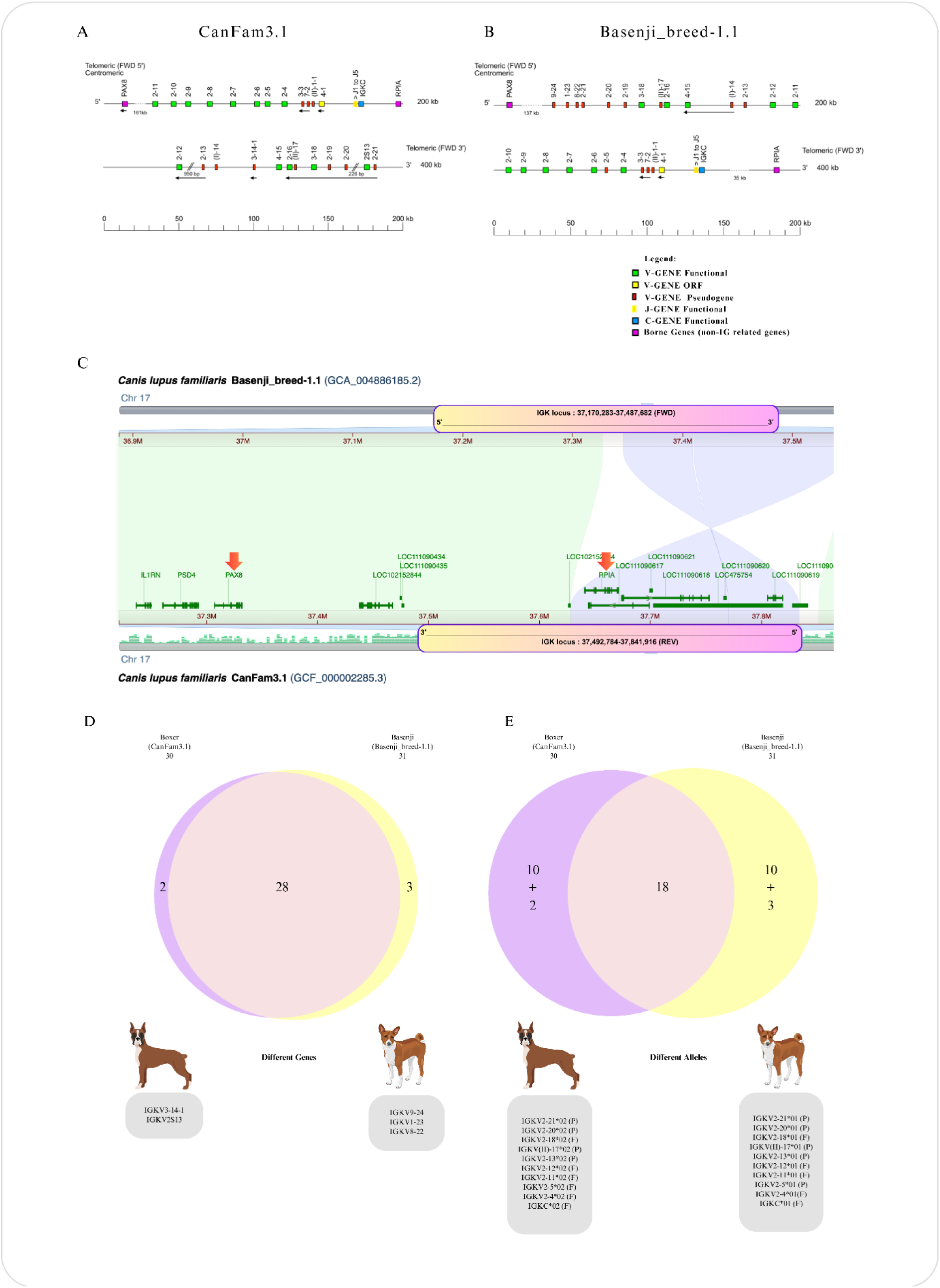
Comparative analysis of the IGK locus in the dog (*Canis lupus familiaris*) IMGT reference genomes: Boxer (CanFam3.1) and Basenji (Basenji_breed-1.1). (A) IMGT reference map of the IGK locus for the Boxer (CanFam3.1) genome, displayed in reverse (REV) orientation, with the presence of two gaps indicated by double slashes. (B) IMGT reference map of the IGK locus for the Basenji (Basenji_breed-1.1) genome in forward (FWD) orientation. Notably, the positions of the V-J-CLUSTER and IMGT 3’ Borne RPIA (Gene ID: 100856455) differ between the two references, with an uncommon pattern observed in the Boxer reference. Reproduction is authorized by IMGT®, the international ImMunoGeneTics information system, and is available on the <u>IMGT® website</u>. (C) Comparative analysis of chromosome 17 between the two IMGT reference assemblies using the Comparative Genome Viewer (CGV). Rectangles delineate the positions of the IGK locus in each assembly on the chromosome. The blue region (hourglass shape) represents alignment sequences in reverse orientation, while white represents non-matched sequences and green denotes matched sequences. Notably, the IMGT 3’ Borne RPIA (Gene ID: 100856455) is mispositioned within the middle of the IGK locus in the Boxer (CanFam3.1) assembly. (D) The Venn diagram illustrates the number of shared IGK genes between the Boxer and Basenji assemblies. (E) Venn diagram illustrating the number of shared IGK alleles between the Boxer and Basenji assemblies. Grey boxes highlight distinct genes (D) and alleles (E) in each assembly, with their respective functional classifications: functional (F) or pseudogene (P). The diagram was generated using Eulerr (https://eulerr.co).

The IGK locus in the Boxer (CanFam3.1) contains 30 genes, comprising 24 V-GENES, 5 J-GENES, and 1 C-GENE, whereas the Basenji (Basenji_breed-1.1) assembly includes 31 genes, totaling 25 V-GENES, with the same number of J-GENES (5) and the C-GENE (1) as in the Boxer (CanFam3.1). The two reference assemblies share the most IGK genes, with 28 out of 33 genes (84.4%) being common, including 22 IGKV, 5 IGKJ, and 1 IGKC gene. However, 5 genes (15.6%) differ between the assemblies: IGKV3-14-1 and IGKV2S13 were found only in the Boxer (CanFam3.1) assembly, while IGKV9-24, IGKV1-23, and IGKV8-22 were present only in the Basenji (Basenji_breed-1.1) assembly, and were located at the (5’ FWD) end of the locus (Figure 6, D and E). Most of the IGKV genes in the Basenji reference assembly (Basenji_breed-1.1) were pseudogenes, whereas in the Boxer reference assembly (CanFam3.1), most IGKV genes were functional (Figure 6, D and E).

#### 2.4.2. Intra-individual IGK locus discrepancies: Boxer case study

Given the discrepancies in IGK locus orientation, V-J-C-CLUSTER position, and gaps in the IMGT reference Boxer (CanFam3.1), the release of a new Boxer assembly (Dog10K_Boxer_Tasha), which is based on sequencing from the same individual as the reference (CanFam3.1), has facilitated improvements and clarify the discrepancies in the annotations of the IGK locus for the first Boxer assembly (CanFam3.1). The locus orientation differs; the Boxer (Dog10k_Boxer_Tasha) has a forward orientation (FWD), contrary to the old reference Boxer (CanFam3.1) (Figure 7, A and B).

**Figure 7.**
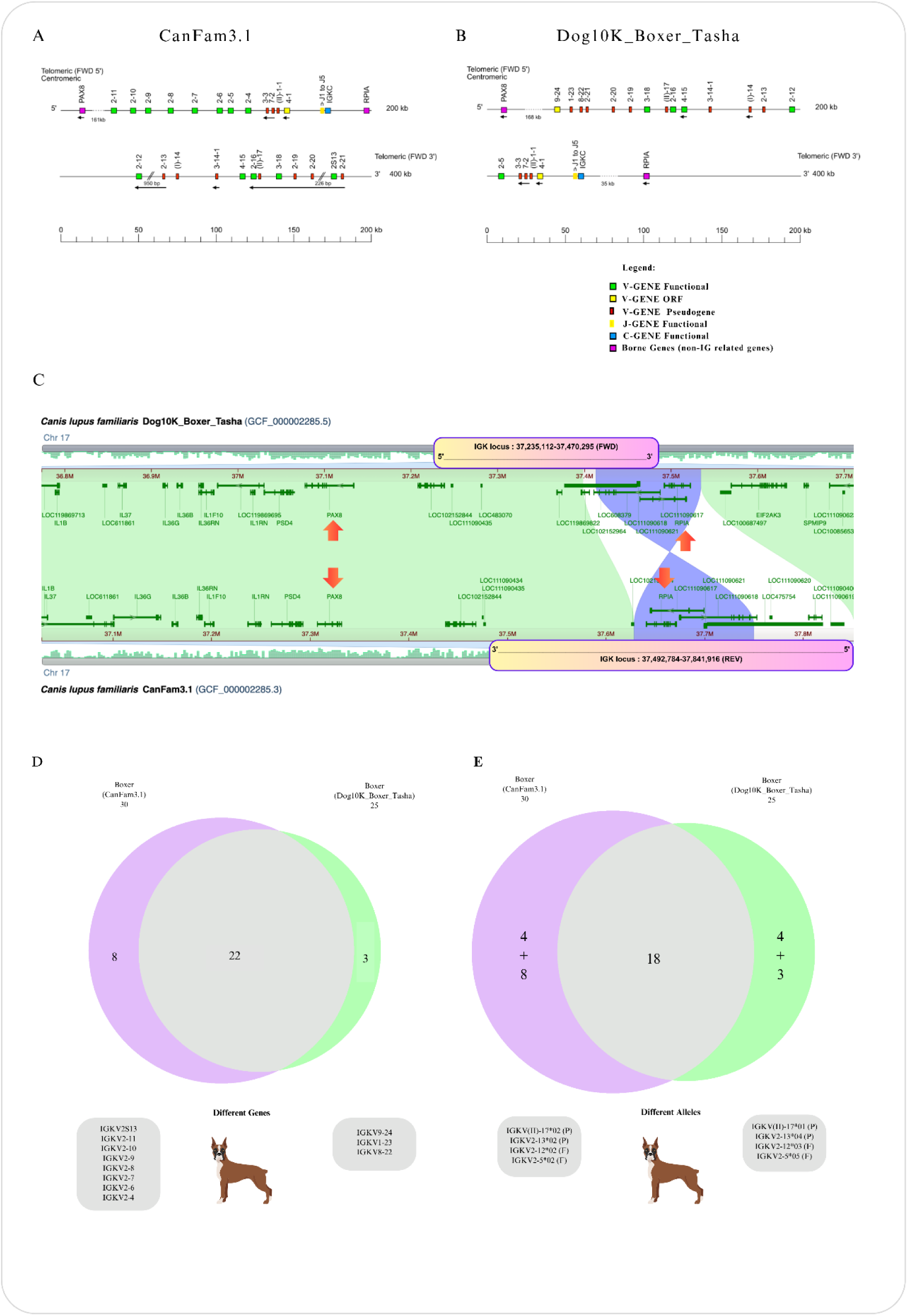
Comparative analysis of the IGK locus in the dog (*Canis lupus familiaris*) IMGT reference genome for the Boxer (CanFam3.1) and the latest version of the Boxer genome (Dog10K_Boxer_Tasha), derived from the same individual but two distinct genome assemblies. (A) IMGT reference map of the IGK locus for the Boxer (CanFam3.1) genome, displayed in reverse (REV) orientation, with the presence of two gaps indicated by double slashes. (B) The IMGT map of the IGK locus for the latest version of the Boxer genome (Dog10K_Boxer_Tasha) in forward (FWD) orientation. Notably, the positions of the V-J-CLUSTER and IMGT 3’ Borne RPIA (Gene ID: 100856455) differ between the two references, with an uncommon pattern observed in the IMGT Boxer reference (CanFam3.1). Reproduction is authorized by IMGT®, the international ImMunoGeneTics information system, and is available on the <u>IMGT® website</u>. (C) Comparative analysis of chromosome 17 between two IMGT Boxer assemblies using the Comparative Genome Viewer (CGV). Rectangles indicate the IGK locus positions in each assembly. The blue hourglass-shaped region represents aligned sequences in reverse orientation, while white areas denote non-matched sequences, and green indicates matched sequences. In the Boxer reference (CanFam3.1), the RPIA gene is located within the IGK locus, whereas in the new Boxer assembly (Dog10k_Boxer_Tasha), RPIA is positioned 35 kb downstream, just after the IGK locus. Additionally, the older assembly (+114 kb) encompasses a larger locus, which may explain the higher number of IGK genes observed in CanFam3.1 compared to Dog10k_Boxer_Tasha. (D) Venn diagram illustrating the number of shared IGK genes between the Boxer assemblies. (E) Venn diagram illustrating the number of shared IGK alleles between the Boxer assemblies. Grey boxes highlight distinct genes (D) and alleles (E) in each assembly, with their respective functional classifications: functional (F) or pseudogene (P). Notably, four genes exhibited allelic polymorphisms between the assemblies. The diagram was generated using Eulerr (https://eulerr.co).

The CGV tool also confirmed that the Boxer assembly (Dog10k_Boxer_Tasha) has the 5’end IMGT flanking gene, “IMGT borne”, PAX8 (PAX8, Gene ID: 701906) at 168 kb upstream of IGKV9-24, the most 5’gene in the locus, and the 3’ end IMGT flanking gene, “IMGT borne”, RPIA (RPIA, Gene ID: 699694) at 36 kb downstream of IGKC (Figure 7, C). The IGK locus length for the Boxer assembly (Dog10k_Boxer_Tasha) spans 235 kb, with no gaps. Interestingly, the IGK locus in the Boxer assembly (Dog10k_Boxer_Tasha) was smaller than the IMGT Boxer reference (CanFam3.1). Both the orientation of the IGK locus and the position of the V-J-C-CLUSTER in the Boxer (Dog10k_Boxer_Tasha) were like the organization observed in the IMGT Basenji reference (Basenji_breed-1.1) (Figure 6, B). This suggests the possibility of annotation errors in the IGK locus of reference Boxer assembly (CanFam3.1) which may have been due to incorrect assembly reconstruction and the gaps in the region of this locus.

The IGK locus of Boxer (Dog10k_Boxer_Tasha) contains 25 genes, comprising 19 V-GENES, 5 J-GENES, and 1 C-GENE. Compared to the Boxer reference (CanFam3.1), the Boxer assembly (Dog10k_Boxer_Tasha) was missing five V-GENES (Figure 7). To determine whether the high gene count in the Boxer reference (CanFam3.1) was due to gene duplications, we employed the Clustal Omega tool, which confirmed the absence of duplicated genes (data not shown). The Boxer assemblies shared 22 out of 33 IGK genes (66.6%), with a discrepancy of 11 genes (33.3%). The eight genes (IGKV2S13, IGKV2-11, IGKV2-10, IGKV2-9, IGKV2-8, IGKV2-7, IGKV2-6, IGKV2-4) were present only in the Boxer reference assembly (CanFam3.1), while three genes (IGKV9-24, IGKV1-23, and IGKV8-22) were present only in the Boxer assembly (Dog10k_Boxer_Tasha) (Figure 7, D and E). Most of the IGKV genes in the new Boxer assembly (Dog10k_Boxer_Tasha) were pseudogenes, whereas in the Boxer reference assembly (CanFam3.1), most IGKV genes were functional. Notably, 4 out of 22 genes (18.2%) shared between the Boxer assemblies (IGKV(II)-17, IGKV2-13, IGKV2-12, and IGKV2-5) exhibit allelic polymorphisms that did not impact their functionality (Supplemental_Table_S2) (Figure 7, D and E).

#### 2.4.3. Nomenclature of IGKV genes

The IGK gene and allele nomenclature were assigned according to IMGT-ONTOLOGY rules, based on the gene position in the locus (NUMBERING). However, this approach is challenging, as allele numbering follows the order of the annotated assemblies, which can result in ambiguities when new genes and alleles are discovered within the same species. After completing the annotation of all assemblies, we found that the new IGKV2-4-1*01 and IGKV2-4-1*02 genes shared the same V-REGION sequences as the new alleles IGKV2-6*02 and IGKV2-9*02, respectively. These new genes (IGKV2-4-1*01, IGKV2-4-1*02) and alleles (IGKV2-6*02, IGKV2-9*02) were first identified in the Bernese Mountain Dog assemblies BD_1.0 and OD_1.0 (Figure 8, A and B). To investigate potential nomenclature errors arising from gene duplication or translocation (possibly caused by assembly errors), we used the Comparative Genome Viewer (CGV) tool (Figure 8, C). The alignment of chromosome 17 between the two Bernese Mountain Dog assemblies revealed discordant regions, indicating a potential translocation in the OD_1.0 assembly and a duplication in the BD_1.0 assembly. A red arrow highlights a possible duplication involving the IGKV2-6*02 allele and the IGKV2-4-1*01 gene in the BD_1.0 assembly, while a blue arrow indicates a potential translocation involving the IGKV2-4-1*02 gene and the IGKV2-9*02 allele in the OD_1.0 assembly. Furthermore, a gap between IGKV2-9*03 and IGKV2-8*03 was identified in the BD_1.0 assembly, complicating confirmation of the translocation and validation of the positioning of IGKV2-4-1*02 (OD_1.0) and IGKV2-9*02 (BD_1.0). Notably, the newly identified genes IGKV2-4-1*01 and IGKV2-4-1*02 were exclusively observed in the Bernese Mountain Dog assemblies. In contrast, the new alleles IGKV2-6*02 and IGKV2-9*02 were shared with the Labrador Retriever assemblies (ROSCfam_1.0 and Yella_v2) and were in their expected genomic positions without evidence of translocations or duplications. Based on these findings, we opted not to alter the existing gene names. The observed assembly inconsistencies highlight the challenges of assigning gene nomenclature (based on position) when assembly errors, such as duplications, translocations, and gaps, are present.

**Figure 8.**
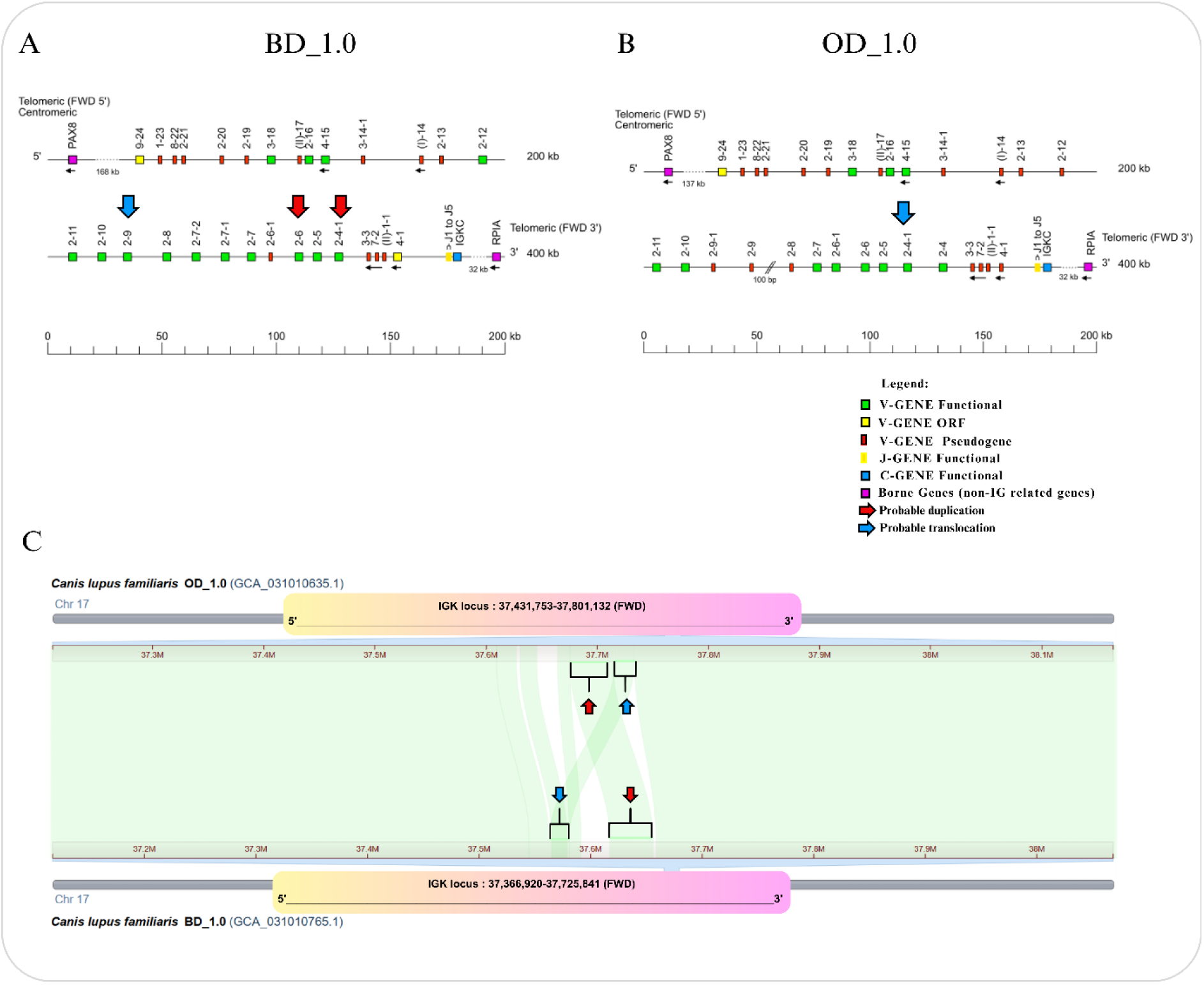
Comparative analysis of the IGK locus between the Bernese Mountain Dog assemblies BD_1.0 and OD_1.0. IMGT maps of the IGK locus for the Bernese Mountain Dog assemblies BD_1.0 (A) and OD_1.0 (B) are presented in forward (FWD) orientation on chromosome 17. Red arrows highlight alleles exhibiting 100% identity in the V-region, suggesting a possible duplication. The blue arrow points to the allele with 100% identity in the V-region shared between the BD_1.0 (A) and OD_1.0 (B) assemblies, indicating a potential translocation. Notably, the OD_1.0 (B) assembly shows a gap (double slash) near the IGKV2-9 allele. Reproduction authorized by IMGT®, the international ImMunoGeneTics information system, available on the <u>IMGT® website</u>. (C) Comparative analysis of chromosome 17 between the two IMGT Bernese Mountain Dog assemblies using the Comparative Genome Viewer (CGV). Rectangles represent the IGK locus positions in each assembly. Red arrows indicate sequence matches of different sizes between the assemblies, suggesting a possible duplication. Blue arrows indicate sequence matches in different locations, implying a possible translocation.

## 3. Discussion

To the best of our knowledge, our study is the first to investigate the IGK gene at the germline level across various dog breeds, offering novel insights into canine immunogenetics. This achievement was made possible by the growing availability of high-quality dog genome assemblies published since 2019, driven by advancements in long-read sequencing technologies such as Oxford Nanopore and PacBio (Halo et al., 2021; Jagannathan et al., 2021; Schall et al., 2023). The comprehensive annotation of multiple assemblies provides several key advantages, including identifying conserved genes and functions, enhanced inter- and intra-breed comparisons, improved annotation accuracy, deeper insight into genetic variability, and the development of a more comprehensive and reliable genomic database.

### The comprehensive annotation of the IGK locus across dog breeds

The comprehensive annotation of the IGK locus across seven dog breeds identified 40 IGK genes, comprising 34 IGKV genes, 5 IGKJ genes, and 1 IGKC gene. In contrast, a previous study by Martin et al. (2018) annotated 19 IGKV genes, 5 IGKJ genes, and 1 IGKC gene, identifying fewer V genes (Martin et al., 2018). The discrepancy may be attributed to differences in methodology, as the authors used RNA-Seq data and based their annotations on rearranged IGK genes rather than genomic sequences.

Our analysis identified seven novel IGK genes, with the majority (five out of seven) located within the Bernese Mountain Dog assemblies (BD_1.0 and OD_1.0) (Schall et al., 2023). These assemblies displayed the longest IGK locus lengths, potentially enabling the discovery of additional genes, consistent with findings that larger locus coverage can uncover greater genomic diversity (Lefranc et al., 2015). However, despite including seven dog breeds, the limited sample size per breed restricted our ability to detect breed-specific genes or deletions, which may reflect the effects of inbreeding or selective pressures. Inbreeding is known to reduce genetic diversity, increase susceptibility to inherited disorders, and elevate genetic load (Axelsson et al., 2021; Jansson & Laikre, 2018). Addressing this limitation, future studies aligned with the Dog10K consortium’s objectives could incorporate larger sample sizes across breeds, enabling more robust detection of breed-specific genomic variations and enhancing our understanding of haplotype structures within canine populations (Meadows et al., 2023).

In dogs, the majority of IGKV genes belong to the IGKV2 subgroup, highlighting its predominance within the canine immune repertoire. Consistent with our findings, Cullen et al. (2022) demonstrated that IGKV2 genes constituted 99.7% of all functional V genes in healthy dogs using 5′ RACE sequencing. Similarly, Martin et al. (2018) reported that 14 out of 19 IGKV genes (73.7%) were classified within the IGKV2 subgroup, reinforcing its dominance. Both studies utilized RNA-Seq to analyze expressed immune repertoires, underscoring the critical role of IGKV2 genes. This subgroup’s predominance may be shaped by evolutionary pressures or specialized immune functions that confer selective advantages. Understanding the dominance of IGKV2 genes could provide valuable insights into breed-specific immune responses, genetic susceptibilities, and opportunities for developing targeted immunotherapies (LeBlanc & Mazcko, 2020).

### Patterns of gene conservation

Among the 40 identified IGK genes, 20 (14 IGKV, 5 IGKJ, and 1 IGKC) were consistently found across all assemblies, reflecting 50% conservation (Figure 2). Notably, 10 of these conserved IGK genes are monomorphic, including IGKV1-23*01, IGKV2-19*01, IGKV4-15*01, IGKV(I)-14*01, IGKV(II)-1-1*01, IGKVJ1*01, IGKVJ2*01, IGKVJ3*01, IGKVJ4*01, and IGKVJ5*01. The monomorphic nature of these genes may indicate fixation within populations, potentially driven by strong selective pressures that favor their retention (Barrett & Hoekstra, 2011). Such pressures might arise from their critical roles in immune function or adaptations to specific environmental challenges. Further exploration of these monomorphic genes could provide insights into the evolutionary dynamics shaping the canine immune repertoire and their implications for disease resistance or susceptibility.

Recent studies of immune repertoire analysis have underscored the predominant usage of specific IGKV-J genes, with five genes accounting for more than 5% of rearrangements: IGKV2-16 (38.7%), IGKV2-5 (20.3%), IGKV2-11 (13.2%), IGKV2-4 (7.6%), and IGKV2-12 (5.7%). Among kappa chain rearrangements, IGKJ1 (33.8%) and IGKJ4 (50.3%) were identified as the most prevalent J gene sets (Cullen et al., 2022), highlighting a highly skewed expressed repertoire. Interestingly, these IGKV genes were not conserved across all annotated assemblies in our study, suggesting potential differences in methodology or underlying genetic diversity. Notably, Cullen et al. relied exclusively on IMGT references for sequence alignment, which may have resulted in a more limited identification of genes. Despite these differences, such findings are pivotal for advancing immune-based therapies and diagnostic tools in veterinary medicine, offering insights into the canine immune system’s structure and function.

### Discovery of new alleles

Through the analysis of nine canine genome assemblies and IMGT references, a total of 97 IGK alleles were identified, comprising 90 IGKV, 5 IGKJ, and 2 IGKC alleles. Notably, 85.3% of the 34 IGKV genes exhibited polymorphisms, highlighting substantial allelic diversity. In contrast, a study by Martin et al. (2018), which analyzed a larger cohort of 107 dogs across 19 breeds, identified a greater number of non-annotated IGK alleles. Their analysis revealed 108 IGK alleles (107 IGKV and 1 IGKC), with the majority (72.8%) maintaining functionality comparable to the reference allele.

Our findings are consistent with those of Martin et al. (2018) and Cullen et al. (2022), both of which reported a predominance of functional IGK genes. In our study, 59 out of 97 alleles were functional. Among the identified IGK genes, 28 (82.35%) exhibited alleles with identical functionality to their reference alleles, while six IGKV genes presented functional variations from their reference alleles. Notably, the genes IGKV2-5 (5 alleles), IGKV2-6-1 (4 alleles), IGKV2-8 (3 alleles), IGKV2-9 (3 alleles), and IGKV4-1 (4 alleles) demonstrated functional diversity. However, the gene IGKV2-4 exhibited the highest degree of polymorphism, with six alleles, all of which were functional. Interestingly, genes exhibiting notable polymorphism were in regions of the locus characterized by “low density,” suggesting potential challenges in assembly or annotation in these areas.

Through Sanger sequencing, we confirmed the *in silico* annotation of two functional *IGKV2* alleles, IGKV2-12*03 and IGKV2-4*04, which had been previously identified in the IMGT/GENE-DB database. The allele IGKV2-12*03 was detected in ten independent canine genome assemblies and was assigned an IMGT confidence score of 3, indicating a high level of reliability in the annotation. The allele IGKV2-4*04, identified in the Cairn Terrier assembly with an IMGT score of 1, also showed full sequence identity with our experimental data. Both alleles were amplified and validated using specific forward primers, confirming their presence and homozygosity in the tested samples.

These results provide experimental evidence supporting the accuracy of the *in silico* annotation and reinforce the existence of functional diversity within the *IGKV2* subgroup. The concordance between the predicted and experimentally validated alleles highlights the reliability of IMGT-based annotations and demonstrates the utility of combining computational and experimental approaches for immunoglobulin gene validation. Moreover, the successful amplification of these alleles from non-lymphoid tissues indicates that such variants are part of the germline repertoire rather than products of somatic rearrangements. Altogether, these findings expand the current understanding of canine immunoglobulin light chain variability and contribute to improving the genomic characterization of the *IGKV* locus across breeds.

### Challenges and limitations

IMGT annotated the dog IGK locus in 2017 using the Boxer genome assembly (CanFam3.1), when no alternative assemblies were available. This annotation, based on orthology with the human IGK locus, which consists of two large, inverted segmental duplications, and was complicated by gaps in the Boxer genome in the IGK region (Debbagh et al., 2024; Engelbrecht et al., 2024). Consequently, the annotation placed half of the IGKV genes upstream and the other half downstream of the IGKJ and IGKC genes, deviating from the typical organization seen in other species (Collins & Watson, 2018). In dogs, a cluster of IGKV genes is found 5′ of a few IGKJ genes, with the IGKJ cluster positioned 5′ of a single IGKC gene. Initially, this atypical structure was interpreted as an evolutionary variation specific to dogs rather than a potential assembly or annotation error, as noted by Collins et al. (2018) and Martin et al. (2018) (Collins & Watson, 2018; Martin et al., 2018).

Given that the Boxer genome (CanFam3.1) has served as the reference for dogs for 13 years and has been cited in over 130 studies, the identification and dissemination of errors in reference genomes and annotations are essential for the scientific community. The publication of the Basenji genome in 2019 (Wallis & Raffan, 2020) and the de novo assembly of the Boxer Tasha genome in 2021 (Jagannathan et al., 2021) provided the opportunity to study and validate the IGK locus orientation in dogs, as discussed in the “Challenges and Limitations” section of this article (Figures 5 and 6).

Interestingly, the de novo Boxer assembly (Dog10K_Boxer_Tasha) could not replace the previous Boxer assembly (CanFam3.1) in the IMGT reference annotations, as it contained five fewer genes than its predecessor. Although the newer Boxer assembly, generated using long-read sequencing technology, demonstrated superior quality, the earlier assembly was based on Sanger sequencing. This discrepancy underscores the importance of annotating multiple genome assemblies to ensure accuracy and reduce errors arising from limitations inherent in single-assembly references (Rhie et al., 2021; Whibley et al., 2021).

Finally, nomenclature issues arose from the IGKV2-4-1*01 and IGKV2-4-1*02 genes, which shared the same V-REGION sequences as the newly identified alleles IGKV2-6*02 and IGKV2-9*02 in the Bernese Mountain Dog assemblies BD_1.0 and OD_1.0. These new genes and alleles will be retained in the IMGT databases and tools; however, we emphasize that if future studies demonstrate the non-existence of any of these variants, they will be removed. In such cases, the nomenclature associated with these genes will not be reused for this locus or species (i.e., non-recyclable names).

### Future perspectives

Enriching the IMGT database with canine sequences from various breeds enables future studies to explore inter and intra-breed differences in the immune repertoire. This study represents the first annotation of the germline sequences of IGK genes in dogs, which provides the advantage of identifying functional and non-expressed genes, such as open reading frames (ORFs) and pseudogenes. These findings are crucial for understanding V-J gene usage and for validating newly identified alleles and genes. Furthermore, the inclusion of alleles in the database will support future research on the characterization of mutations in the immunoglobulin genes of the IGK locus, which is of critical importance in the prognosis of chronic lymphocytic leukemia (CLL) as shown by the European Research Initiative on CLL (ERIC) (Chatzikonstantinou et al., 2024). However, this area has not been sufficiently investigated in dogs due to the absence of alleles from various breeds (Avery, 2020; Rout et al., 2018).

Additionally, our research identified specific challenges in canine genome sequencing, particularly in regions with low gene frequency (low density). It could be attributed to complex repetitive regions and high heterozygosity of Immunoglobulin loci, as observed in humans, presenting significant challenges for current sequencing methods (Rodriguez et al., 2023). A promising approach to addressing the low coverage in these regions is the use of long-read sequencing technologies combined with specialized pipelines. For instance, IMGT/StatAssembly was designed to evaluate the accuracy and completeness of genome assemblies in complex regions, providing a valuable framework for overcoming these challenges.

In conclusion, our findings reveal substantial allelic variability within the canine IGK locus. Through Sanger sequencing, we experimentally validated two newly annotated IGKV2 alleles, both previously identified in silico and classified as functional. This allelic diversity represents an important factor to consider in the development of immunologically based therapeutic strategies, including vaccines, immunotherapies, and monoclonal antibody treatments for cancer and other immune-related diseases.

## 4. Methods

The annotation of the IGK locus was conducted according to the IMGT biocuration pipeline, as previously described (Pégorier et al., 2020). The *Canis lupus familiaris* genomic sequences from new assemblies were analyzed and annotated according to the IMGT dog reference sequences^1^ [^1^https://www.imgt.org/IMGTrepertoire/index.php?section=LocusGenes&repertoire=locusAssembly&species=dog&group=IGK).] from Boxer (CanFam3.1) and Basenji (Basenji_breed-1.1) (Martin et al., 2018).

### 4.1. Criteria for the selection of novel dog assemblies

The canine assemblies were selected by searching for “*Canis lupus familiaris*” in the NCBI Datasets in December 2023 (Kitts et al., 2016). The selection criteria were defined by IMGT® on chromosomal-level organization, haploid assemblies, and complete genome representation. Thus, nine assemblies from seven different breeds were chosen for the quality of their IGK locus, which fulfilled the standard IMGT criteria for assembly selection^2^ [^2^https://www.imgt.org/IMGTScientificChart/Assemblies/IMGTassemblyquality.php]: “Basenji” (UNSW_CanFamBas_1.2, GenBank Assembly ID: GCA_013276365.2), “Bernese Mountain Dog” (BD_1.0 and OD_1.0, GenBank Assembly ID: GCA_031010765.1 and GCA_031010635.1), “Boxer” (Dog10K_Boxer_Tasha, GenBank Assembly ID: GCA_000002285.4), “Cairn Terrier” (CA611_1.0, GenBank Assembly ID: GCA_031010295.1), “German Shepherd” (ASM864105v1, GenBank Assembly ID: GCA_008641055.1), “Great Dane” (UMICH_Zoey_3.1, GenBank Assembly ID: GCA_005444595.1), and “Labrador Retriever” (ASM1204501v1 and ROS_Cfam_1.0, GenBank Assembly ID: GCA_012045015.1 and GCA_014441545.1).

### 4.2. IGK locus extraction

The IGK locus on chromosome 17 for each assembly was localized and extracted using BLAST (v. 2.15.0) by comparing sequences against IGK reference sequences from Basenji (Basenji_breed-1.1) and Boxer (CanFam3.1) in IMGT/GENE-DB (v. 3.1.37), corresponding to IMGT/LIGM-DB accession numbers IMGT000067 and IMGT000002, respectively (Giudicelli et al., 2005). The locus position and orientation were defined based on the flanking non-IG genes, known as “IMGT bornes,” identified as paired box 8 (PAX8, Gene ID: 701906) and ribose 5-phosphate isomerase A (RPIA, Gene ID: 699694). The IMGT bornes provide a standardized method for comparing IG and TR locus delimitation across species (Lefranc & Lefranc, 2022). Distances between the IMGT bornes and the flanking IG genes (IGKV9-24 upstream and IGKC downstream) were measured. If the borne genes were absent or distances exceeded 10 kb, an extraction interval was set to extend 10 kb upstream of IGKV9-24 and 10 kb downstream of IGKC.

### 4.3. Quality evaluation of the IGK locus and integration in IMGT

The extracted IGK locus sequence was evaluated by vector contamination to detect foreign DNAs such as vector, linker, adapter, and primer regions using VecScreen (Schäffer et al., 2018). The Comparative Genome Viewer (CGV) (Rangwala et al., 2024) from NCBI was employed to analyze the IGK locus between two assemblies, allowing for the comparison of locus orientation, and the investigation of potential deletions, insertions, translocations, and inversions that could affect IGK locus annotation. The default parameters included a minimum alignment size of 10 kb, with adjustments to the viewer to display non-best-placed alignments and show both forward and reverse alignments. The IMGT/LIGMotif internal tool was used to evaluate the orientation of genes in the locus, the total number of genes by group and subgroup, and to find and delimit the gaps when they were present (Lane et al., 2010). After the quality evaluation of the locus, the corresponding nucleotide sequences were extracted from the NCBI chromosome sequences, and entries were created in IMGT/LIGM-DB with an IMGT accession number (IMGT’ followed by 6 digits) (Giudicelli et al., 2005).

### 4.4. V, J, and C genes annotation

The V, J, and C genes were identified and delineated along the IMGT/LIGM-DB genomic sequences of the IGK locus using IMGT/LIGMotif (Giudicelli et al., 2005; Lane et al., 2010). IGK genes were characterized and classified through BLAST alignments, Clustal Omega, and the application of IMGT’s unique numbering and annotation rules according to the IMGT Scientific Chart (Johnson et al., 2008; Lefranc et al., 2003; Sievers et al., 2011). The nomenclature for *Canis lupus familiaris* IGK genes follows IMGT-ONTOLOGY^3^ [^3^https://www.imgt.org/IMGTScientificChart/Nomenclature/IMGTnomenclature.html] concepts for gene and allele functionality (Giudicelli & Lefranc, 2012). In agreement, the nomenclature for IGK genes of Labrador Retriever assembly (ROS_Cfam_1.0) was approved by the Vertebrate Gene Nomenclature Committee (VGNC) in 2024 (Jones et al., 2023). The NCBI Third Party Annotation (TPA)^4^ [^4^https://www.ncbi.nlm.nih.gov/genbank/tpa/] accession numbers were assigned to two assemblies from the Bernese Mountain Dog breed: BD_1.0 and OD_1.0, corresponding to IMGT/LIGM-DB accession numbers BK066061 and BK066062, respectively. The whole data of IGK genes were integrated into IMGT/GENE-DB (Giudicelli et al., 2005) and the biocuration data for genes, alleles, and proteins were entered into IMGT web resources^5^ [^5^https://www.imgt.org/IMGTrepertoire/].

## 5. Primer Design, PCR Amplification, and Sequence Analysis

Seven forward primers and one consensus reverse primer were designed to validate alleles belonging to the *IGKV2* subgroup, considering the V-EXON region, which presented the highest number of alleles. The primer sequences are listed in Table 5. Additional information can be assessed in Supplemental Table S4.

**Table 5.**
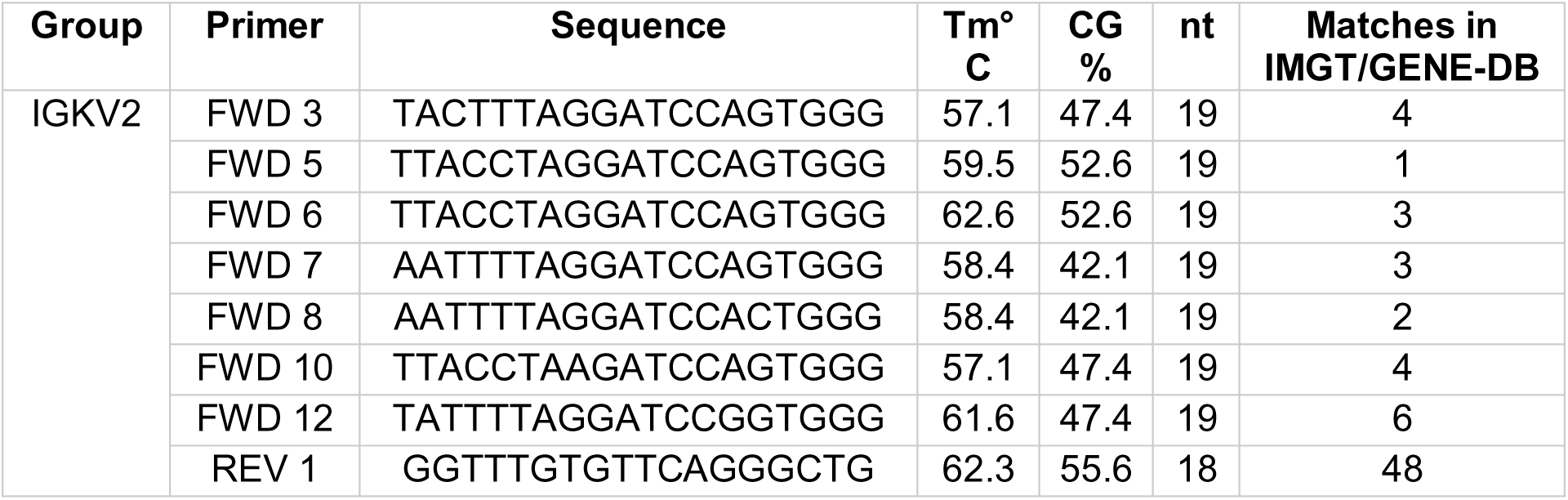
Primers for validation of *Canis lupus familiaris* IGKV2 alleles.

Genomic DNA (gDNA) was extracted from non-lymphoid cardiac tissue samples of three dogs (two females and one male) obtained from the LOCT Laboratory Biobank (Ethics approval: CEUA 7602220224). Each PCR reaction contained 100 ng of gDNA, 200 nM of each primer, 1× Platinum™ PCR Master Mix (Invitrogen, USA), and molecular biology-grade water in a final volume of 50 µL.

PCR amplification was performed on a C1000 Touch™ Thermal Cycler (Bio-Rad, USA) under the following conditions: initial denaturation at 94 °C for 2 min; followed by 25 cycles of denaturation at 94 °C for 30 s, annealing at 55 °C for 30 s, and extension at 72 °C for 30 s.

Amplicons were visualized on a 2% agarose gel and purified using the QIAquick PCR Purification Kit (Qiagen, Germany) according to the manufacturer’s instructions. Sanger sequencing was performed on an ABI 3730 DNA Analyzer (Applied Biosystems, Thermo Fisher Scientific, USA) using the BigDye™ Terminator v3.1 Cycle Sequencing Kit.

The obtained sequences were analyzed with FastQC (v0.12.1) to assess the PHRED quality scores. Reads were trimmed using Sickle (v1.33; https://github.com/najoshi/sickle) with default Sanger parameters. Forward and reverse sequences were then analyzed using IMGT/V-QUEST (version 3.7.0; reference directory 202531-1) and compared with the IMGT reference database. Ambiguous nucleotides were resolved using overlapping information from the complementary sequence, when applicable.

## 6. Data access

IGK locus sequences and all related annotations from 9 dog assemblies are available in the IMGT/LIGM database (ref) with the accession numbers: BK066061, BK066062, IMGT000179, IMGT000180, IMGT000181, IMGT000182, IMGT000183, IMGT000184, and IMGT000185. The annotations are also available in the TPA database on the NCBI for BK066061 and BK066062. IGK genes are managed and distributed through IMGT/GENE-DB and the IMGT reference directory for the analysis of expressed repertoire with IMGT/V-QUEST and IMGT/HighV-QUEST is available at https://www.imgt.org/IMGTindex/directory.php. Additionally, the original contributions presented in the study are included in Supplemental_Table_S3.

## 8. Acknowledgements

We would like to express our heartfelt gratitude to the whole IMGT® team for their expertise and incredible collaboration in carrying out this study. We are also grateful to Prof. Marie-Paule Lefranc for her invaluable contribution to building the dog’s IGK reference annotations on IMGT. We also extend special thanks to Dr. Marina Omelchenko, responsible for the Comparative Genome Viewer (CGV) tool at NCBI, for her support in assisting us and providing comparisons between the canine genomes. We extend our thanks to Dr. Ruth Seal and her team from the Vertebrate Gene Nomenclature Committee (VGNC) for recognizing the nomenclature of the annotated genes and making it accessible in their database. Their efforts have significantly supported the standardization and dissemination of genetic information. We thank the São Paulo Research Foundation (FAPESP), Grants number 23/05951-5 and 21/09982-7, and the Brazilian Coordination for the Improvement of Higher Education Personnel (CAPES) for the financial support of the TKG Ph.D. thesis and internship at IMGT®. IMGT® is a member of the French Infrastructure Institut Français de Bioinformatique (IFB) as well as a member of BioCampus, MAbImprove, and IBiSA.

## 9. Author contribution

TKG, HF, and SK conceived the study. TKG analyzed the data and drafted the manuscript. AP, MG, VG, JJM, GF, PG, and GZ analyzed the data, validated the results, and supported the updates to the databases. TKG, TGLA, and JCCL performed the wet lab assay, PCR, and Sanger sequencing. VG and GF developed and updated the internal and external IMGT® tools for data analysis. SK, HF, VG, GF, JJM, and AK discussed and reviewed the manuscript. SK supervised the findings and the write-up of this work. All authors contributed to the article and approved the submitted version.

